# HRS dephosphorylation at membrane contact sites promotes sorting within multivesicular endosomes

**DOI:** 10.64898/2026.03.10.710128

**Authors:** Minoo Razi, Louise H. Wong, Douglas Grimes, Thomas Burgoyne, Pedro Fale, Ewan MacDonald, Emily R. Eden, Michael J. Clague, Sylvie Urbe, Clare E. Futter

## Abstract

HRS is a receptor tyrosine kinase substrate and component of the ESCRT machinery. It enables sorting of ubiquitylated cargo into multivesicular endosomes (MVEs) for lysosomal degradation but also functions in receptor recycling. We show that co-depletion of its ESCRT-0 binding partners, STAM1 and STAM2, recapitulates defects in EGF receptor (EGFR) sorting onto intraluminal vesicles (ILVs) but does not mirror the increased MVE size evident after HRS depletion. Using mutagenesis of the endogenous gene or introduction of APEX2-tagged HRS variants, we find that HRS Y329/334 phosphorylation is dispensable for EGFR sorting and MVE size control. AnnexinA1 mediates endosomal contact with the ER and promotes EGF-stimulated ILV formation. We show that reduced ILV formation on AnnexinA1 depletion is accompanied by increased HRS phosphorylation, likely through reduced HRS dephosphorylation by ER-localised PTP1B. Conversely, Y329/334 mutation renders ILV formation insensitive to AnnexinA1 depletion. Our data suggest that rapid HRS dephosphorylation at ER:MVE contacts promotes efficient ILV formation.

## Introduction

HRS (also referred to as HGS) is part of ESCRT0, which forms the first of a series of protein complexes (ESCRT0-III) that sort endocytosed ubiquitylated proteins like EGF-stimulated EGFR from the limiting membrane of multivesicular endosomes (MVEs) onto the intraluminal vesicles (ILVs) that accumulate in the MVE lumen^1,2^. This redirects the receptors from the recycling pathway so that when the MVE fuses with the lysosome, the receptors are degraded. HRS is recruited to endosomes by binding to phosphatidylinositol-3 phosphate (PtdIns3*P*) through its FYVE domain and to ubiquitylated endocytic cargo through its ubiquitin-interacting motif (UIM), a property shared with its ESCRT0 binding partner, STAM1 or STAM2^3–6^. HRS recruits clathrin to endosomes to form a coat that scaffolds HRS in domains on the limiting membrane to concentrate ubiquitylated cargo, as well as regulating ILV size and shape^6–10^. HRS also recruits the ESCRTI component, TSG101, as part of the process of transfer of ubiquitylated cargo to later ESCRTs^11–13^. Ordered polymerisation of ESCRTIII drives inward invagination of the limiting membrane, with scission coupled to ESCRT dissociation catalysed by the AAA ATPase, VPS4^2^. HRS is also implicated in endocytic recycling to the plasma membrane^14,15^ and TGN^16^ and egress of LDL-derived cholesterol from the endocytic pathway^17^, roles proposed to be independent of downstream elements of the ESCRT machinery. The regulation of HRS activity to fulfil these different roles remains to be elucidated and we specifically aimed to investigate the hitherto poorly characterised role of reversible HRS tyrosine phosphorylation.

HRS is one of the most prominently tyrosine phosphorylated substrates following EGF stimulation^18,19^, with the major phosphorylation sites downstream of the EGFR being Y329 and Y334^5^ (Phosphosite plus www.phosphosite.org). Inhibitor studies indicate that a non-receptor tyrosine kinase of the Src family downstream of the EGFR is responsible for this phosphorylation^5,20^. EGF-stimulated HRS phosphorylation is highly dependent on receptor internalisation^4,19^ and on association of HRS with endosomes since deletion of either the FYVE domain or the UIM domain abrogates HRS phosphorylation^4,5^. Although HRS requires endosome association for phosphorylation to occur, most phosphorylated HRS is found in the cytosol, suggesting that after phosphorylation at least a proportion of HRS is released from the endosomal membrane^4,5^. Consistently, a transient loss of HRS from endosomes has been demonstrated that is co-incident with and dependent upon EGF-stimulated HRS phosphorylation^21^. Live cell imaging of tagged ESCRT components has revealed waves of ESCRT recruitment and dissociation controlled by clathrin^10^. Surprisingly, EGF stimulation modestly increased dwell time of HRS on endosomes that contained EGF^22^, raising the question of how this can be reconciled with phosphorylation-induced release from the endosomal membrane. HRS can also be monoubiquitylated in a UIM-dependent manner, which prevents binding to ubiquitylated cargo^23^ ^24^ and HRS phosphorylation has been linked with its ubiquitylation and degradation^25^.

Despite its prominence, the function of EGF-stimulated HRS phosphorylation has been surprisingly difficult to resolve. EGF stimulation upregulates ESCRT-dependent ILV formation^26^ and EGF-stimulated HRS phosphorylation has been reported to promote EGFR degradation^21,25^, suggesting that EGF-stimulated HRS phosphorylation could promote ESCRT-dependent HRS activity.

HRS phosphorylation peaks transiently about 8 minutes after EGF stimulation^4,21,27^, suggesting that it is subject to dephosphorylation by phosphatases. We and others have found that HRS can be dephosphorylated by the protein tyrosine phosphatase, PTP1B^27,28^. We previously showed that EGFR is dephosphorylated by PTP1B at membrane contact sites that form between EGFR-containing MVEs and the ER^27^. In principle, the cytosolic pool of phosphorylated HRS could interact with PTP1B independently of membrane contact sites. However, it is also possible that membrane contacts between endosomes and the ER are sites of interaction of PTP1B with components of the ESCRT machinery to allow local control of ESCRT activity. PTP1B depletion not only enhances HRS phosphorylation but also inhibits ILV formation^27^, supporting a model in which timely HRS dephosphorylation regulates ILV formation. This hypothesis is in keeping with the timing of EGF-stimulated ILV formation which continues for at least 30 minutes after EGF stimulation^10,29^, in marked contrast to the transient HRS phosphorylation. Membrane contact sites between EGFR-containing MVEs and the ER are tethered by AnnexinA1:S100A11 complexes^30^. Abrogation of these contacts by AnnexinA1 depletion inhibits EGF-stimulated ILV formation^26^, suggesting a role for the contacts in regulating ILV formation.

We aimed to determine the role of HRS phosphorylation/dephosphorylation in regulating its function. Our data point to a suppression of ESCRT-dependent ILV formation by transient HRS phosphorylation that is relieved by HRS dephosphorylation at AnnexinA1-dependent contact sites between MVEs and the ER. These contacts are subject to regulation by ESCRT0, which may co-ordinate contact site assembly/disassembly to allow ILV formation to occur.

## Materials and methods

### Cell culture and transfection

All cell lines, Human hTERT-RPE1 (ATCC, CRL-4000) and HeLa (ATCC, CCL2) were maintained in Dulbecco’s Modified Eagle’s Medium (DMEM) supplemented with 10% FBS and 100 units/mL penicillin, 100□µg/mL streptomycin (Invitrogen). HeLa S3 Flp-In host cells were a gift from Ian Prior, (University of Liverpool).

#### CRISPR knock-in cells

The HGS gene from hTERT-RPE1 cells was sequenced and a guide sequence targeting HGS was selected using Benchling and analysed for potential off-target sites. An HDR template was designed, replacing A with T in the codons corresponding to Y329 and Y334, replacing these amino acids for phenylalanine. The guide, HDR template and Alt-R™ S.p. Cas9 Nuclease V3 were ordered from Integrated DNA Technologies (Leuven, Belgium). After 18 hrs of growth, individual cells were grown in a 96 well plate, and resulting colonies were expanded and selected by PCR amplification using primers that amplify the edited region. Induced mutations in the targeted region were confirmed by Sanger sequencing.

The guide homology sequence was: 5’-AGCUCGCACGGUAUCUCAAC

The HDR template sequence was: 5’GGG AGT GAC CCC CTC ATT GCC TGC AGC TCG CAC GGT TTC TCA ACC GCA ACT TCT GGG

#### Flp-In cells

PCR products of APEX2-HRS, GFP-HRS and YYFF mutants were amplified from pEGFP-C1-HRS and YYFF constructs encoding mouse HRS^5^ and cloned into a pEF5/FRT/V5 TOPO vector to generate Flp-In compatible vectors. To generate stable cell lines, HeLa S3 Flp-In host cells were transfected with each of the generated plasmids together with pOG44 (Flp-recombinase) at a ratio of 1: 9. Following hygromycin B selection (150-200 µg/ml), single colonies were isolated, amplified and tested for transgene expression, verified by Immunofluorescence and western blotting.

For siRNA knockdown, cells were transfected by using either Lipofectamine RNAiMAX reagent (Invitrogen) or by the nucleofector (Lonza) following the manufacturer’s instructions.

### Antibodies

EGFR (sc-373746); GAPDH (sc-47724); STAM2 (sc-365600) Tubulin (sc-32293); from Santa Cruz. Hrs (ab155539) and STAM1 (ab155527) from abcam. Annexin1A (610066) from BD Transduction Labs; Tyr1068 Phospho-EGF receptor (3777) from Cell Signalling. Anti-phosphospecific Y334-HRS was previously described^5^.

### siRNAs

The non-targeting control siRNA was SilencerTM Select negative control No. 1 (Themo Fisher Scientific, Life Tech), On-targetPlus siRNA against STAM1(J-011423-05) and STAM2 (J-017361-06) Annexin1A (J-011161-07) was from Dharmacon. siRNA targeting HRS used in APEX2-mHRS HeLa cells was AGA GAC AAG UGG AGG UAA A dTdT. siRNAs targeting HRS used in GFP-mHRS HeLa cells (Supplementary Figure 3) were HRS-1, 5′-CGUCUUUCCAGAAUUCAAA-3′ and HRS-2, 5′-UGGAAUCUGUGGUAAAGAA-3′).

### Western blotting

Cells were lysed in ice-cold TNTE buffer [20□mM Tris-HCl pH7.4, 150□mM NaCl, 5□mM EDTA, 1% Triton-X100, complete protease inhibitor cocktail (Roche 04693124001), PhosSTOP (Roche 04906837001)] except for Western blots shown in Supplementary Figure 3. For the latter, cells were lysed in RIPA buffer (10mM Tris-HCl pH 7.5., 150mM NaCl, 1% Triton X-100, 0.1% SDS, 1% Sodium deoxycholate supplemented with mammalian protease inhibitors (Sigma) and PhosSTOP inhibitors (Roche). Lysates were cleared by centrifugation and resolved on NuPAGE Bis-Tris 4%–12% gels (Life Technologies) under reducing conditions, followed by transfer onto PVDF (Amersham Hybond p 0.2micron) or nitrocellulose membranes.

Following incubation with relevant primary and secondary antibodies the blots were developed by ECL prime western blotting reagent. Blots were imaged using Bio-Rad Chemidoc. Densitometry was performed with ImageJ.

### Electron Microscopy

Cells were treated as described previously^31^. Briefly, cells were serum starved for 1 h prior to stimulation with 100 ng/ml EGF (Sigma) with gold-conjugated anti-EGFR antibody in DMEM for the specified time. After fixation cells were embedded as described previously^32^.

Before embedding, cells expressing APEX2-mHRS were incubated with 1.5 mg/ml DAB (TAAB) in TRIS buffer supplemented with 0.02% H_2_O_2_ for 30 min in the dark following fixation as described^33^. Sections were viewed using a Joel 1400 plus TEM and either; Deben NanoSprint12 Camera with AMT software or Gatan Orius CCD Camera with Digital Micrograph software. Micrographs were analysed using ImageJ.

### Electron tomography

Tilt series were collected over a range of ±60° from 150nm thick serial sections on a JEOL 1400 plus TEM fitted with a Gatan Orius SC1000B camera and using SerialEM (University of Colorado, US). Dual axis tomograms were generated and joined together using IMOD (University of Colorado, US). Microscope Imaging Browser (University of Helsinki, Finland) and IMOD were used to segment the data and ChimeraX (University of California, San Francisco, US) used to visualise the resulting model (stretched along the z-axis by 1.95 to correct for the anisotropic resolution).

### Fluorescence microscopy

For results shown in Figure 2, cells were serum-starved for an hour before stimulation with 100 ng/ml Alexa Fluor-conjugated EGF (E35350 or E13345, ThermoFisher Scientific), fixed in 4% paraformaldehyde and images collected using a Leica Stellaris 5; HC PL APO 63x/1 Objective and Hamamatsu Flash camera. For results shown in Supplementary Figure 3, cells were serum starved for 16 hours before stimulation with 20 ng/ml EGF, fixed as above and stained with anti-EGFR (R1 from CRUK) and then AF594-coupled donkey anti-mouse (Invitrogen). Images were captured using a Marianas spinning disk confocal microscope (3i) using a 63× 1.4 NA Zeiss Plan Apochromat lens, FLASH4 sCMOS (Hamamatsu) camera.

## Results

### Distinguishing between ESCRT-dependent and ESCRT-independent roles of HRS by STAM1/2 depletion

We compared the effects of HRS depletion with depletion of the alternate ESCRT0 partners, STAM 1 and 2, reasoning that only those features caused by loss of ESCRT-dependent HRS functions would be mimicked by STAM depletion. siRNA targeting either HRS or STAM1/2 reduced the amount of both STAM and HRS, indicating that reduction of individual ESCRT0 components reduces the stability of other components of the complex (Supplementary Figure 1), in keeping with a previous demonstration that HRS suppresses STAM degradation^34^. However, more HRS can be depleted by HRS-targeting siRNA than by STAM-directed siRNA. This suggests that there is a pool of HRS that can be depleted by siRNA targeting HRS but not by siRNA targeting STAM1/2, likely representing an HRS pool that operates independently of STAM. Consistent with this hypothesis, the effects of HRS and STAM1/2 depletion could be uncoupled. siRNA-treated cells were incubated with EGF for 25-30 minutes in the presence of an antibody directed to the extracellular domain of EGFR coupled to colloidal gold (EGFR-gold). We have previously shown that this EGFR-gold allows the normal trafficking route of the endocytosed EGFR to be followed by transmission electron microscopy (TEM)^29^. As with previous studies^26,29,31^, MVEs were defined as electron lucent vacuoles containing one or more discrete ILVs and monodisperse gold. These can be readily distinguished from lysosomes, which are electron dense, frequently contain membrane whorls and contain gold that has aggregated in the acidic degradative lysosomal lumen. As expected, depletion of HRS in HeLa cells inhibited sorting of EGFR onto ILVs and was accompanied by a substantial increase in mean MVE cross-sectional area (Figure 1A-C). In STAM1/2-depleted cells sorting of EGFR onto ILVs was inhibited to an extent not significantly different to that of HRS depletion, consistent with this being an ESCRT-dependent role of HRS (Figure 1A-B). However, MVE size upon STAM1/2 depletion was not significantly different to that of control cells (Figure 1A and C), indicating that endosome enlargement is caused primarily by loss of ESCRT-independent HRS function(s). We previously found that depletion of HRS generates unusually small ILVs in a subset of MVEs and proposed that this was due to a loss of ESCRT-dependent ILV formation leading to upregulation of ESCRT-independent ILVs^35^. Consistent with this hypothesis we found that STAM1/2 depletion, like HRS depletion, led to a decrease in mean ILV size (Figure 1D).

**Figure 1.**
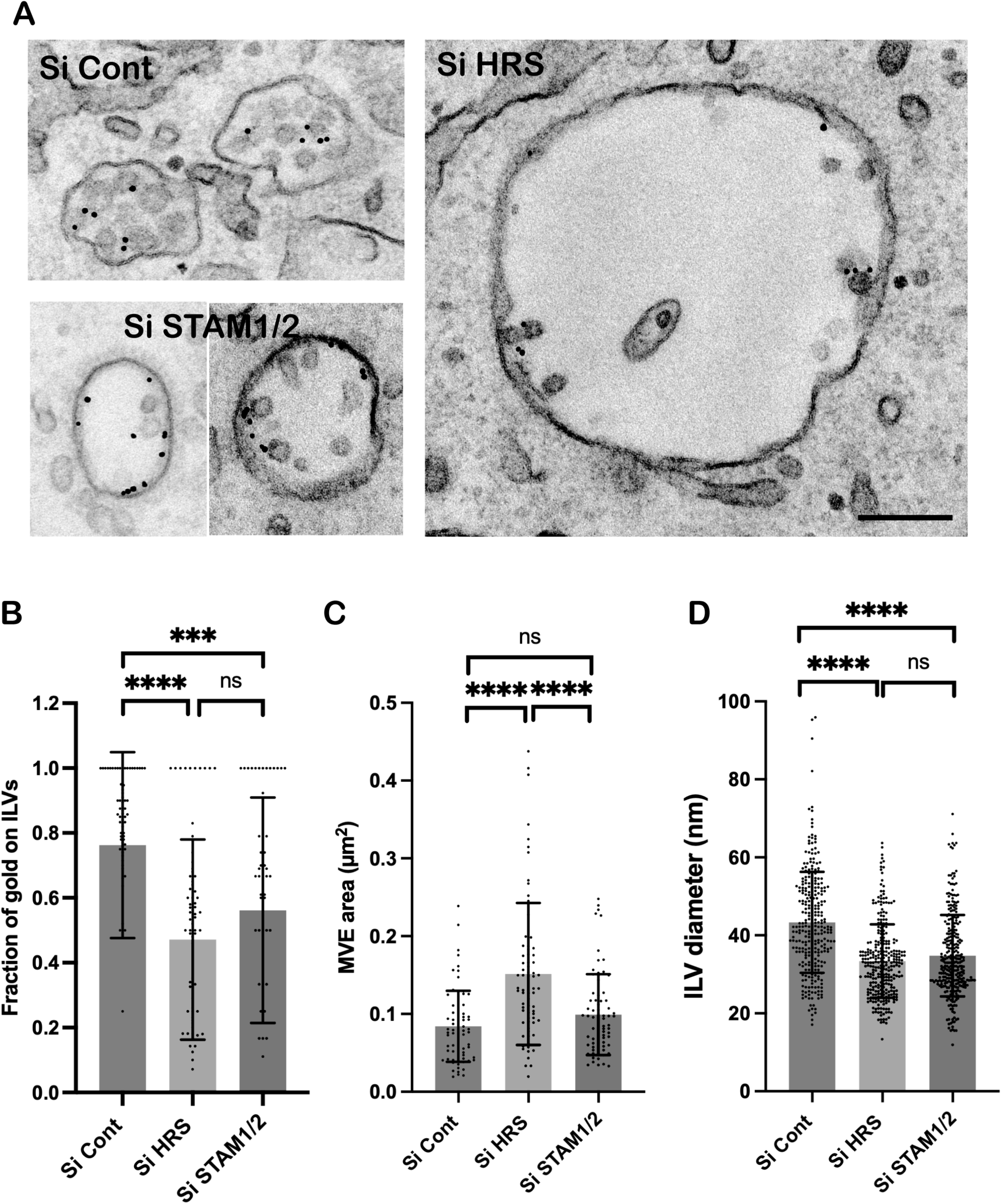
Comparison of effects of HRS and STAM1/2 knockdown on MVEs in HeLa cells. (A) TEM micrographs of HeLa cells treated with control siRNA (Si Cont) or siRNA targeting HRS or both STAMs 1 and 2 and stimulated with EGF and anti-EGFR gold for 25 minutes before fixation. (B) Fraction of gold on ILVs, (C) MVE area, and (D) ILV size quantified in random sections. Results are Mean +/-SD of at least 100 MVEs from 3 separate experiments and analysed by unpaired t-test. ****p<0.0001, ***p<0.004. Scale bar=200nm.

### Mutation of HRS phosphorylation sites does not affect EGFR endocytosis or degradation

Independent cell models were generated to investigate the role of phosphorylation of the two major EGF-stimulated HRS phosphorylation sites, Y329 and Y334. Overexpression of HRS has previously been shown to inhibit EGFR sorting and cause endosomal enlargement^5,36,37^. CRISPR/Cas9 mutagenesis was used to replace tyrosines 329 and 334 with phenylalanines in endogenous HRS in hTERT-RPE1 cells and two knock-in cell lines homozygous for the mutations were isolated. In HeLa cells, the ‘Flp-In’ system was used to express either wild-type or Y329/334F mutant mHRS fused to the peroxidase, APEX2, or GFP. Depletion of the endogenous orthologue with siRNA specifically targeting the human protein allows the effect of near endogenous levels of the tagged mouse orthologue to be investigated in this model (Supplementary Figure 2A).

Western blotting with a specific anti-phospho Y334 (PY334) HRS antibody showed a peak of phosphorylation 8 minutes after EGF stimulation in the parent hTERT-RPE1 cell line (Supplementary Figure 2B) but no detectable signal in the two Y329/334F CRISPR cell lines (Figure 2A and Supplementary Figure 2B). Similarly, expression of APEX2-mHRS-Y329/334F in HeLa cells depleted of endogenous HRS prevented the HRS phosphorylation seen in cells expressing wild-type APEX2-mHRS 8 minutes after EGF stimulation (Figure 2C). After stimulation with EGF, the distribution of endocytosed EGF (Figure 2B and 2D) or EGFR (Supplementary Figure 3) was indistinguishable between cells expressing wild-type and Y329/334F HRS, indicating no major effect of HRS phosphorylation deficiency on EGFR endocytosis. Stimulation of wild-type and HRS phosphorylation deficient cells with EGF and measurement of EGFR levels by Western blotting revealed no clear differences in the EGFR degradation rate between the cell lines expressing wild-type and phosphorylation-deficient HRS (Supplementary Figure 2B and 3). This was in contrast to previously published data showing expression of phosphorylation-deficient HRS considerably reduced EGFR degradation after 2 and 4 hours of EGF stimulation^21^. As that study also used cycloheximide to prevent new EGFR synthesis, the effects of phosphosite-mutation on EGFR levels after EGF stimulation in the presence of cycloheximide was investigated. There was a substantial reduction in EGFR levels following 2 and 4 hours of EGF stimulation that was indistinguishable between wild-type and mutant cells (Figure 2E).

**Figure 2.**
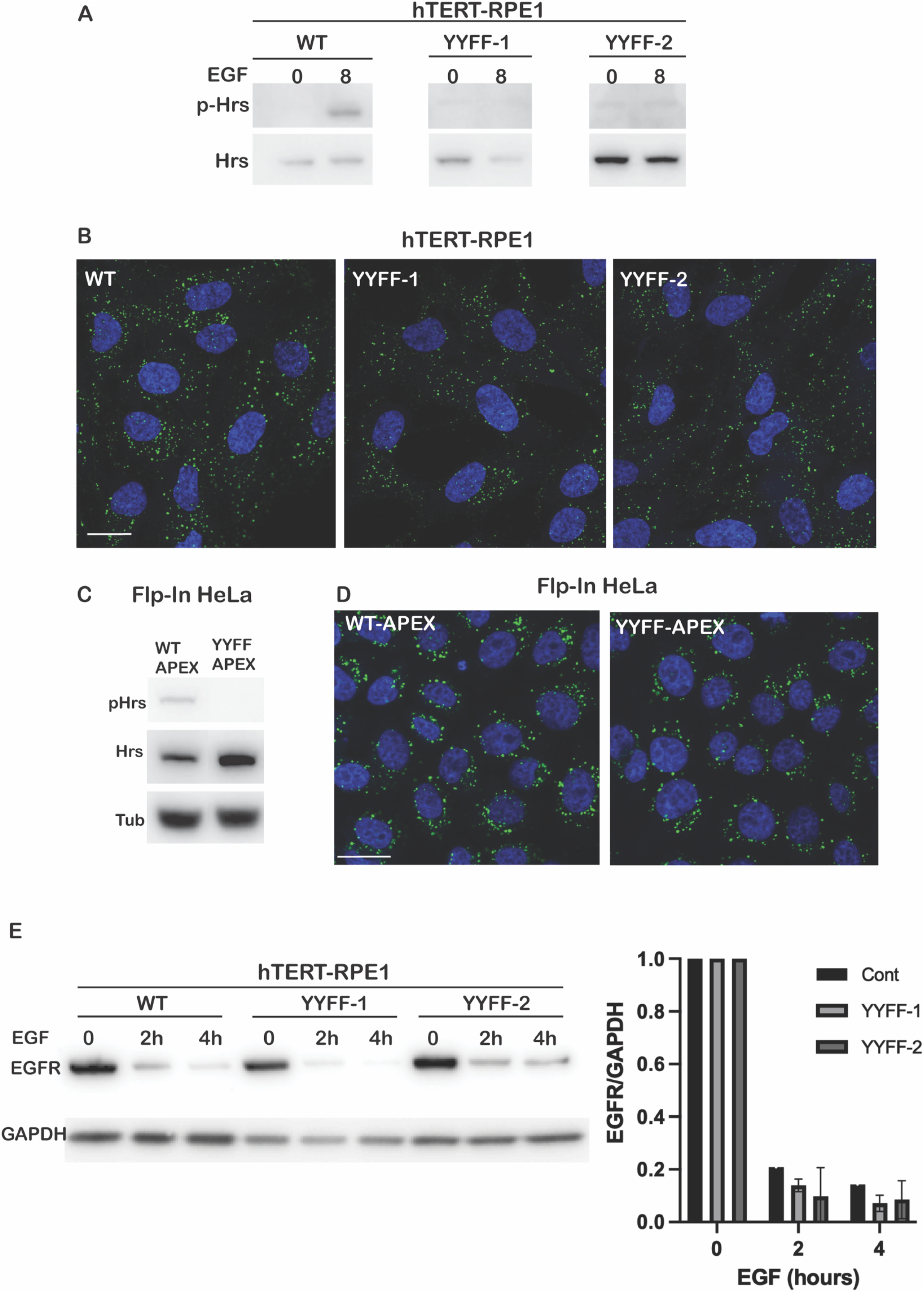
Characterisation of independent models expressing HRS-Y329/334F. Two clones of hTERT-RPE1 cells in which Y329 and Y334 in endogenous HRS has been replaced with phenylalanine using CRISPR (A and B) and Flp-In HeLa cells expressing wild-type APEX2-mHRS or APEX2-mHRS-Y329/334F (C and D) were generated. (A and C) phospho-HRS (Y334) and total Hrs analysis by Western blot of protein extracts from (A) hTERT-RPE1 and (B) HeLa models with endogenous HRS depleted stimulated with EGF for 8 mins. (B and D) Confocal images of hTERT-RPE1 (B) and HeLa models with endogenous HRS depleted (D) incubated with fluorescent EGF (green) for 30 minutes at 37°C. Single confocal slices at the level of the nucleus are shown. Scale bar=20 µm. (E) EGFR quantification by western blot in hTERT-RPE1 models treated with EGF and cycloheximide (CHX) for 0-4 hours. Results are Mean +/− SD of 3 observations.

**Figure 3.**
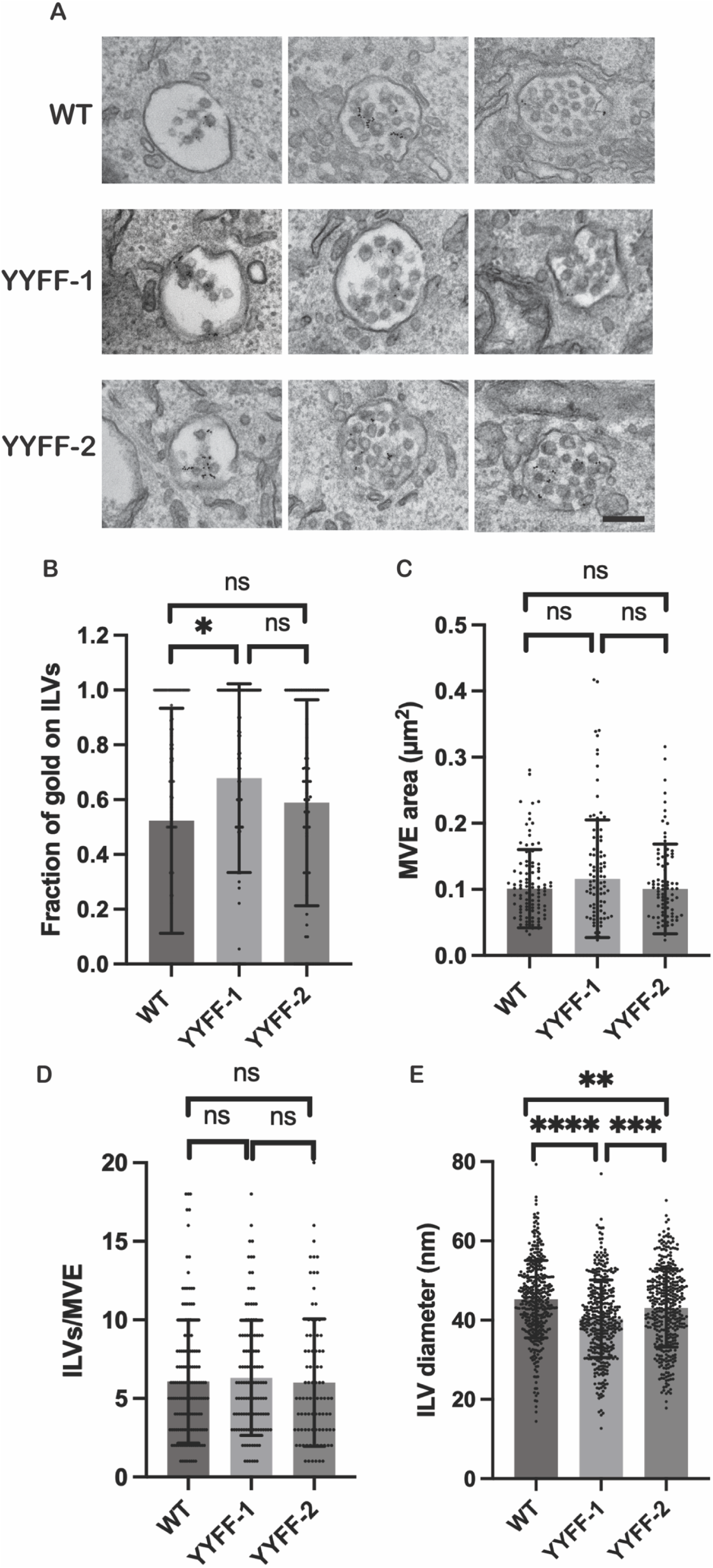
Effects of HRS-Y329/334F mutation on sorting within MVEs in hTERT-RPE cells. (A) TEM micrographs of hTERT-RPE1 parent and two clones of Y329/334F stimulated with EGF and anti-EGFR gold for 25 mins before fixation. (B) Fraction of gold particles on ILVs, (C) MVE area, (D) number of ILVs/MVE and (E) ILV size quantified in random sections. Results are Mean +/-SD of at least 100 MVEs from 3 separate experiments and analysed by unpaired t-test. ****p<0.0001, ***p<0.004. Scale bar=200nm.

### Mutation of HRS phosphorylation sites has little effect on MVE size, EGFR sorting onto ILVs, or ILV formation

To determine the effects of HRS phosphorylation on sorting within MVEs, cells were incubated with EGF and EGFR-gold for 25-30 minutes since at this time, ILV formation is well advanced but there is little lysosomal delivery. As shown in Figure 3A MVEs in Y329/334F CRISPR cells were morphologically indistinguishable from those in the parent cell line. Quantitative analysis revealed a small increase in sorting of EGFR onto ILVs in one phosphorylation-deficient cell line, but this was not reproduced in the other mutant line (Figure 3B). MVE size and number of ILVs per MVE showed no significant differences between the parent and HRS phosphorylation-deficient lines (Figure 3C and 3D). Both Y329/334F lines showed a small decrease in mean ILV size (Figure 3E), although this was less than observed following depletion of total Hrs (see Figure 1D) and was significantly different between the two mutant cell lines. A similar quantitative analysis was performed on the HeLa cell model expressing APEX2-mHRS with endogenous HRS depleted. There were no clear morphological differences between MVEs containing wild-type or phosphorylation deficient APEX2-mHRS (Figure 4A). There was no significant difference in the sorting of EGFR onto ILVs or ILV number per MVE between cells expressing wild-type or phosphorylation deficient APEX2-mHRS (Figure 4B and C). Furthermore, there was no detectable difference in ILV size (Supplementary Figure 4). There was a small increase in MVE size in cells expressing APEX2-mHRS-Y329/334F (Figure 4D) but this was not reproduced in the CRISPR HRS-Y329/334F cell lines. Our data across all cell models indicated that mutation of the Y329/334 phosphorylation sites of HRS left ESCRT-dependent and ESCRT-independent HRS functions intact.

**Figure 4.**
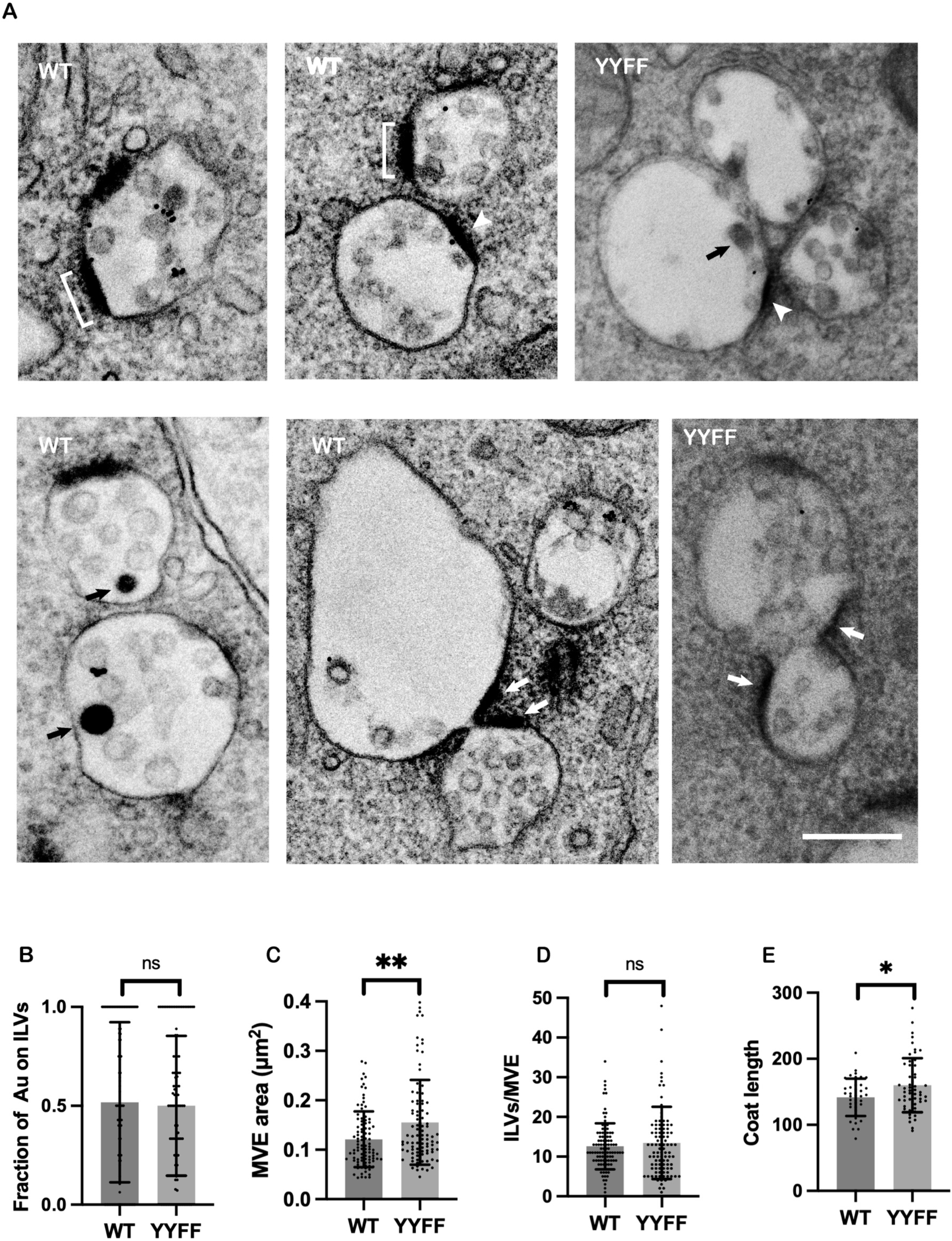
APEX2-mHRS provides an independent model to measure effects of HRS-Y329/334F on sorting within MVEs and to visualise the HRS-containing coat. (A) TEM micrographs of Flp-In HeLa cells expressing wild-type APEX2-mHRS or APEX2-mHRS-Y329/334F both depleted of endogenous HRS. Cells were stimulated with EGF and anti-EGFR gold for 25 mins before fixation and processing for DAB histochemistry. Brackets indicate examples of electron dense DAB positive APEX2-Hrs positive coats. White arrows indicate DAB positive regions adjacent to the neck of hourglass profiles, white arrowheads indicate DAB positive regions adjacent to contacts between MVEs and black arrows indicate DAB positive ILVs. (B) Fraction of gold on ILVs, (C) number of ILVs/MV, (D) MVE area and (E) length of each discrete coat quantified in random sections. Results are Mean +/-SD of at least 100 MVEs from 3 separate experiments and analysed by unpaired t-test. **p= 0.0016, *p=0.013. Scale bar=200nm.

### APEX2 tagging allows visualisation of HRS-containing coated domains on MVEs

The ascorbate peroxidase activity of the APEX2 tag, in combination with DAB histochemistry, allows high resolution localization of the tagged protein^38^ with a greater sensitivity and improved structural preservation compared to immuno-EM techniques. This technique also readily lends itself to electron tomography and consequent visualization of the 3D arrangement of the coat. APEX2-mHRS was present in discrete regions on flattened portions of the MVE limiting membrane (Figure 4A). These were most electron dense adjacent to the limiting membrane and frequently had a less dense ‘fluffy’ appearance on the cytoplasmic side. The APEX2 tag allows the HRS-positive relative coat length to be measured with an accuracy previously not possible. The mean coat length derived from random sections of cells treated for 8 minutes with EGF (the time of peak HRS phosphorylation) was moderately increased in cells expressing phosphorylation deficient HRS (Figure 4E) consistent with phosphorylation promoting release of wild-type HRS from the limiting membrane. The APEX2 tag revealed further interesting features of the HRS-containing coat that were not markedly affected by mutation of phosphor-sites. These included rare but striking DAB positive ILVs (black arrows in Figure 4A) and HRS-positive coats immediately adjacent to membrane contacts between nneighbouring MVEs (white arrowheads in Figure 4A). Additionally, HRS-positive domains were frequently found next to the necks of hourglass profiles of MVEs (white arrows in Figure 4A). Conventional EM suggested the presence of multiple discrete coats on single MVEs but this does not preclude the presence of a connection between coated domains in another section plane, as indicated by the tomographic reconstruction shown in Figure 5A (and Supplementary Videos 1 and 2). In this example what appear to be two separate coated domains in single tomographic slices are actually joined, as shown in the 3D model (Figure 5B in burnt orange). The model also allows comparison of the size of the HRS-containing coated domain with the size of ILVs (in gray), with the coated domain being much larger than the surface area of ILVs.

**Figure 5.**
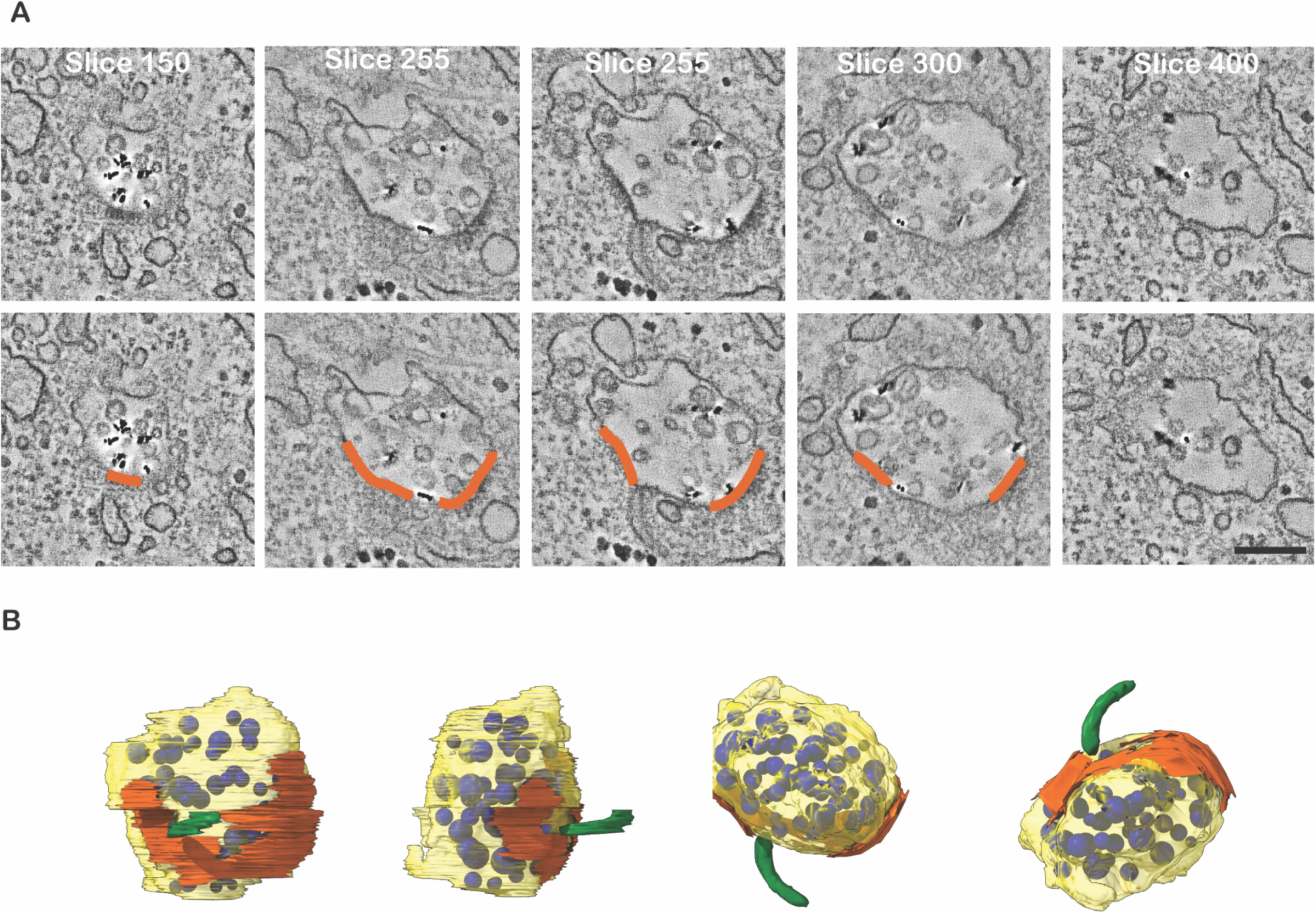
Electron tomography allows 3D reconstruction of HRS positive coated domains on the MVE limiting membrane. Flp-In HeLa cells expressing wild-type APEX2-mHRS were depleted of endogenous HRS before stimulation with EGF and anti-EGFR gold for 25 mins and processed for TEM. Serial section electron tomographic data was generated. (A) Slices of the resulting tomogram with coats highlighted in orange. Full data stack shown in Supplementary Video 1. (B) Model showing segmented MVE limiting membrane (yellow), ILVs (blue), ER (green), and APEX2-mHRS-containing coat (burnt orange). Rotating view shown in Supplementary Video 2. Scale bar=200nm.

### AnnexinA1 depletion enhances HRS phosphorylation

As HRS lacking the major EGF-stimulated phosphorylation sites can support ILV formation, we wondered whether dephosphorylation of HRS has a regulatory role in this process. HRS can be dephosphorylated by PTP1B and we have shown that PTP1B interacts with endosomal substrates like EGFR at membrane contact sites tethered by AnnexinA1:S100A11 complexes^30^. To determine whether AnnexinA1-dependent membrane contact sites regulate HRS phosphorylation, the effects of AnnexinA1 depletion on HRS phosphorylation in wild-type cells was determined. As shown in Figure 6A-D, Western blotting cell lysates of control siRNA-treated cells with anti-phospho-Y334-HRS showed a peak of phosphorylation 8 minutes after EGF stimulation followed by a decline such that phosphorylation was barely detectable after 30 minutes. Depletion of AnnexinA1 significantly enhanced HRS phosphorylation in both hTERT RPE cells (Figure 6C) and HeLa cells (Figure 6D) after 8 minutes EGF stimulation but did not prevent the subsequent loss of HRS phosphorylation.

**Figure 6.**
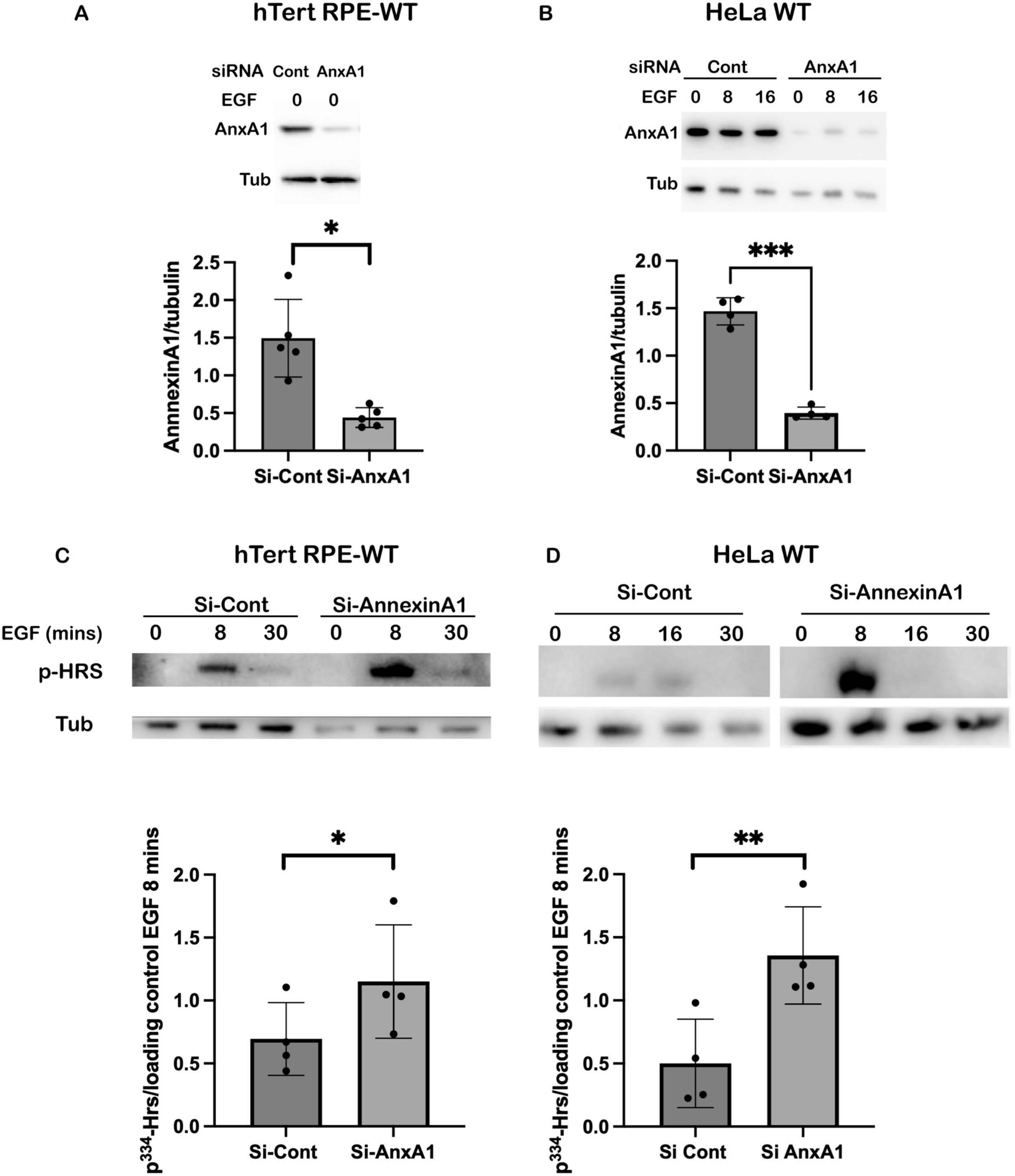
Effects of AnnexinA1 depletion on HRS phosphorylation state. hTERT-RPE cells expressing wild-type HRS (A and C) and wild-type HeLa cells (B and D) were treated with control si-RNA or si-RNA targeting AnnexinA1. (C and D) Cells were incubated with EGF for the indicated times and blotted with anti-phospho-Y334-HRS and anti-tubulin antibodies. Ratio of phosphorylated HRS (pHRS) to tubulin for cells stimulated with EGF for 8 minutes are shown. Results are Mean +/-SD of 3 experiments and analysed by paired t-test. ***p=0.0004, **p=0.0032, *p<0.015.

### Phosphorylation-deficient HRS is resistant to the effects of AnnexinA1 depletion on ILV formation

The enhanced HRS phosphorylation at early time points after EGF stimulation upon Annexin A1 depletion could be a consequence of reduced membrane contact sites and hence reduced dephosphorylation by PTP1B. AnnexinA1 depletion also inhibits sequestration of EGFR on the ILVs of MVEs^26^ and the longer residence time on the limiting membrane of the MVE could lead to enhanced HRS phosphorylation. If dephosphorylation of HRS is required for efficient ILV formation we would expect cells expressing the Y329/334F-HRS to be resistant to the effects of AnnexinA1 depletion. Consistently, as shown in Figure 7A-C, in wild-type cells AnnexinA1 depletion caused a reduction in ILV formation, as we have previously shown in multiple cell types^26^. In contrast, ILV formation in phosphorylation deficient cells was unaffected by AnnexinA1 depletion. Taken together, our data indicate that AnnexinA1 supports ILV formation by suppressing HRS phosphorylation.

**Figure 7.**
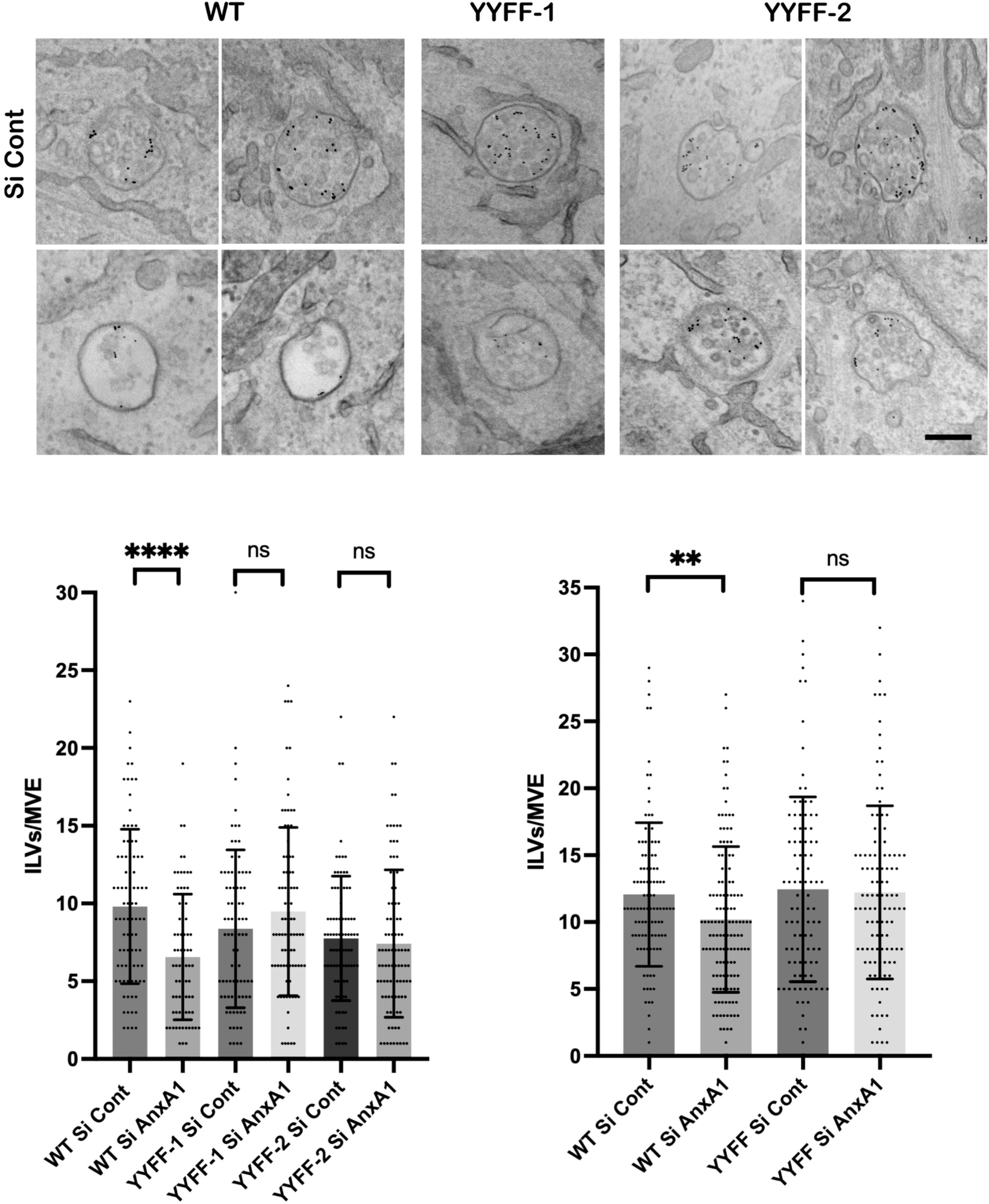
Effects of AnnexinA1 depletion on ILV formation in phosphorylation-deficient cells. hTERT-RPE cells (A and B) and Flp-In HeLa cells (C) expressing wild-type or Y329/334F HRS were treated with control si-RNA or si-RNA targeting AnnexinA1. HeLa cells were also depleted of endogenous HRS. Cells were incubated with EGF and anti-EGFR-gold for 30 mins before fixation. TEM micrographs of hTERT-RPE cells are shown in A. Number of ILVs/MVE in hTERT-RPE cells (B) and Flp-In HeLa cells (C) were quantified in random sections. Results are Means +/− SD of at least 100 MVEs from 3 separate experiments and analysed by unpaired t test. ****p<0.0001, **p=0.0048. Bar=200nm.

### HRS and STAM1/2 depletion but not HRS phosphosite mutation inhibit membrane contact site formation/maintenance

The enhanced phosphorylation of HRS following AnnexinA1 depletion is consistent with HRS being dephosphorylated at MVE:ER contact sites, raising the question of the relationship between ESCRT-mediated ILV formation and membrane contact sites. The presence of an HRS-containing clathrin coated domain would likely preclude contact site formation but membrane contact sites have been observed at the edge of clathrin-coated domains^39^. APEX2-tagged HRS and DAB histochemistry offer a unique opportunity to visualise the relationship between the HRS-containing coat and contact sites at the TEM level. Figure 8A shows examples where the DAB positive coat sits immediately adjacent to contact sites (contacts indicated by arrows) between MVEs and the ER. Also shown is an example where the membranes of the the ER and MVE are separated by the DAB positive coat (indicated by arrowheads). In the tomogram shown in Figure 5 an ER tubule (green) can be observed in contact with the MVE limiting membrane in a gap between two coated areas. If the HRS-containing coat precludes the formation of a membrane contact but Hrs is dephosphorylated at membrane contacts it is likely that either the coat or the contact must be disassembled to allow formation of the other. One way to co-ordinate this process would be for ESCRT0 to play a role in membrane contact site formation/disassembly in addition to its established role in clathrin coat recruitment. We therefore investigated the effect of HRS or STAM1/2 depletion on membrane contact sites between the ER and EGFR-containing MVEs.

**Figure 8.**
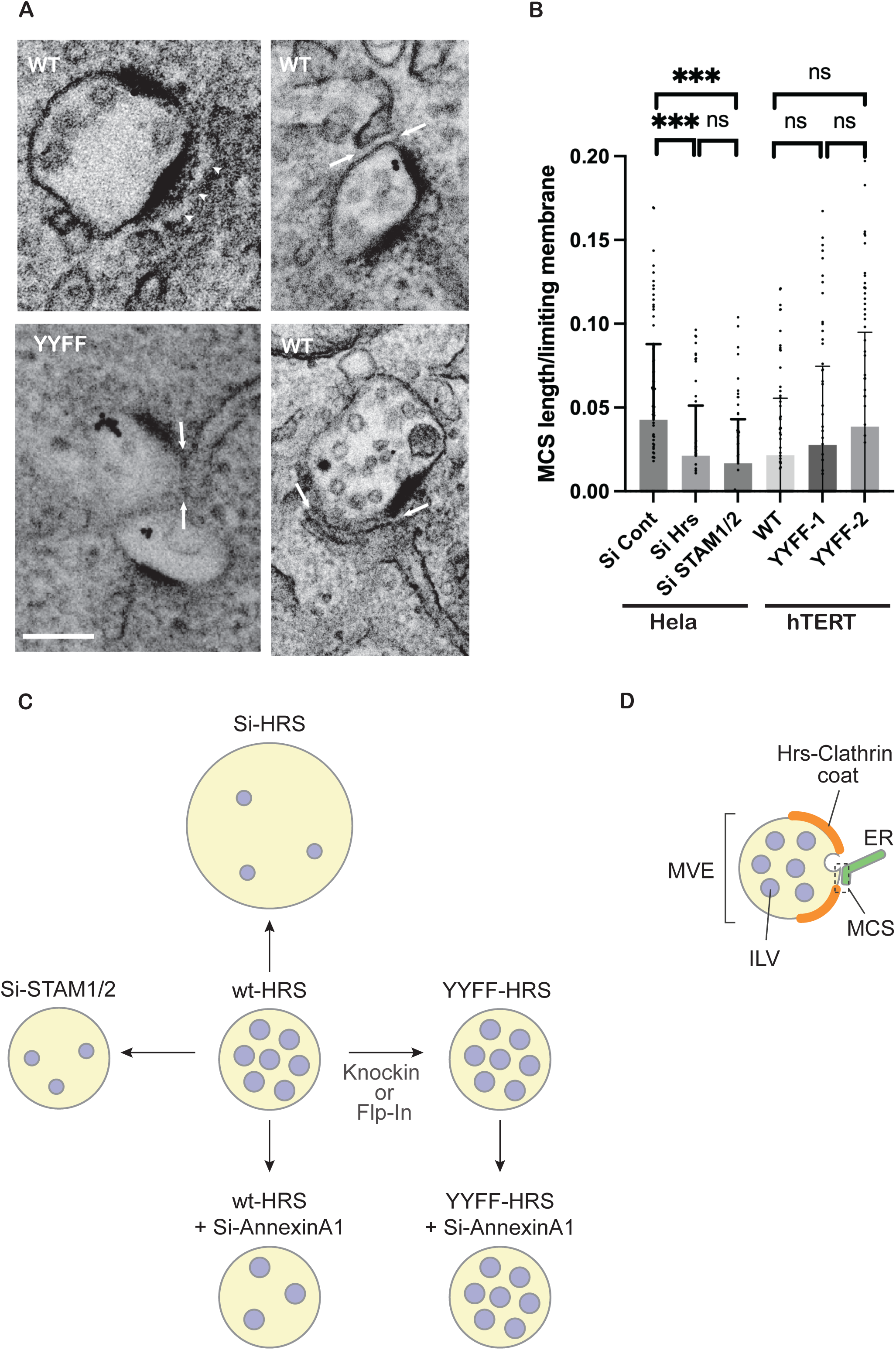
Role of ESCRT0 in regulating membrane contact sites between MVEs and the ER. (A) TEM micrographs of Flp-In HeLa cells expressing wild-type or Y329/334F APEX2-mHRS with endogenous HRS depleted and stimulated with EGF and anti-EGFR-gold for 25 mins. Arrows indicate membrane contacts between MVEs and the ER. Arrrowheads indicate an example where the coat is between the MVE limiting membrane and the ER, precluding contact formation. (B) Quantification of membrane contact site length in micrographs of HeLa cells treated with control siRNA or siRNAs targeting HRS or STAMs 1/2 and hTERT-RPE cells expressing wild-type or Y329/334F HRS stimulated with EGF and anti-EGFR-gold for 30 mins before processing for TEM. Results are Means +/− SD of at least 100 MVEs from 3 separate experiments and analysed by unpaired t test. ***p<0.001. Bar=200nm. (C) Enlarged MVEs are a feature of HRS depletion but not STAM depletion or Y329/334F mutation, indicating that increased MVE size reflects loss of ESCRT- and phosphorylation-independent HRS function(s). Reduced cargo sorting onto ILVs/ILV size on HRS depletion was mirrored by STAM depletion but not Y329/334F mutation, indicating loss of ESCRT-dependent, phosphorylation-independent HRS function(s). Effects of Y329/334 mutation were only revealed upon AnnexinA1 depletion which inhibited ILV formation in wild-type but not Y329/334F-HRS mutant cells. (D) A possible model of the relationship between the HRS-containing clathrin coat, membrane contact site and ILV formation. HRS becomes phosphorylated within the scaffold of the clathrin coat. At the edge of the coat, membrane contact sites form allowing dephosphorylation of HRS (by ER-localised PTP1B), which retains HRS (and STAM) on the MVE limiting membrane to recruit later ESCRTs and promote ILV formation.

HRS and STAM depletion reduced the length of ER:MVE membrane contact site/length of limiting membrane by approximately 50%, indicating a role for ESCRT0 in contact site formation and/or maintenance (Figure 8B). The effects of AnnexinA1 depletion strongly suggest that these contacts have a role in regulating HRS phosphorylation. To determine whether the reverse is also true, that HRS phosphorylation state regulates contact sites, these were quantified in hTERT-RPE cells expressing wild type versus phosphorylation-deficient HRS (Figure 8B). Contact site lengths were variable but there was a trend towards increased membrane contact site length in phosphorylation deficient HRS expressing cells. This suggests that HRS phosphorylation is not a requirement for contact site formation or maintenance but could have a subtle role in regulating their stability

## Discussion

Comparing the effects of HRS with STAMs1/2 depletion enabled the distinction of ESCRT-dependent and -independent roles of HRS and allowed us to show that both ESCRT-dependent sorting within MVEs and ESCRT-independent control of MVE size proceeded in the absence of Y329/334-HRS phosphorylation (Figure 8C). Enhancing Y329/334 phosphorylation through AnnexinA1 depletion was accompanied by reduced ILV formation in wild-type but not Y329/334 HRS-expressing cells (Figure 8C), suggesting that the hyperphosphorylated HRS induced by AnnexinA1 depletion in wild-type cells, but absent from Y329/334 HRS-expressing cells, was inhibitory to ILV formation.

### Loss of ESCRT-independent functions contributes to endosome enlargement in HRS-depleted cells

We previously concluded that inhibition of ILV formation was insufficient to generate the highly enlarged endocytic vacuoles observed in HRS-depleted cells^31^. Here we show that depletion of HRS or its alternate binding partners in ESCRT0, STAM 1 and 2, induce a similar reduction in sorting of EGFR onto ILVs and in ILV size but STAMs1 and 2 depletion does not reproduce the vacuolar enlargement induced by HRS depletion, despite reducing HRS levels. Thus, endosome enlargement is largely caused by loss of ESCRT-independent HRS functions, presumably mediated by the pool of HRS that is resistant to STAM depletion. HRS has been implicated in recycling of a G protein coupled receptor^14^, the TrkB receptor^40^, in WASH recruitment and associated actin-dependent recycling of MT1-MMP and unstimulated EGFR to the plasma-membrane^15^, as well M6PR and Shiga Toxin B recycling to the TGN^15,16^. Reduced membrane recycling could contribute to the endosomal enlargement in HRS-depleted cells.

### APEX2-tagged mHRS allows high resolution imaging of endosomal coats in 3D

The role of HRS in ESCRT-mediated sorting of ubiquitinated cargo into domains of the limiting membrane of MVEs is well established^31,41^. However, the relationship between these domains and ILV formation remains controversial. One study concluded that clathrin recruited by HRS defines the timing of ILV formation, as well as ILV size and shape, such that one coat would result in one ILV^10^. Another concluded that multiple ILVs could bud from a single coat^22^. Our data with APEX2-mHRS allowed the visualisation of the HRS-coated domain in 3D which showed that the size of the coat could considerably exceed the surface area of an ILV and suggests that there is not a direct relationship between coat and ILV size.

### HRS Y329/334 phosphorylation is not necessary for efficient EGFR degradation or ILV formation

Previous studies have concluded that HRS Y329/334 phosphorylation is necessary for efficient EGFR degradation^21,25^ whilst we found no impact of the same mutations on kinetics of EGFR degradation in independent models. The reasons for this discrepancy are not entirely clear. CRISPR – mediated mutation of endogenous HRS phosphorylation sites overcomes potential confounding issues of HRS overexpression or small amounts of endogenous wild-type protein remaining in HRS-depleted cells. Multiple tyrosine phosphorylation sites have been identified in HRS with Y132 and 216 in the VHS and FYVE domains respectively, phosphorylated downstream of the colony stimulating factor receptor and fibroblast growth factor receptor 1 (see Phosphosite Plus www.phosphosite.org). Whilst Y329 and 334 are the major phosphorylation sites downstream of the EGFR^5^, the phosphorylation of alternative sites following mutation of Y329/334 that could compensate for loss of Y329/334 cannot be excluded. However, the lack of an effect of HRS phosphorylation deficiency on EGFR degradation is in keeping with our comprehensive analysis of EGFR sorting at the level of the MVE, which showed no effect on sorting of EGFR onto ILVs or ILV formation 25-30 mins after EGF stimulation. Although in this study we focused on HRS phosphorylation, STAM is also phosphorylated upon EGF stimulation^20,28^. A previous study found that phosphorylation-deficient STAM could support EGFR degradation, but led to prolonged phosphorylation specifically of AKT, suggesting a role for STAM phosphorylation in regulating specific signalling pathways downstream of the EGFR^28^. The distinct patterns of HRS and STAM tyrosine phosphorylation after stimulation by different growth factor receptors are also consistent with a role for ESCRT0 phosphorylation in regulating the diversity of signalling outputs^20^.

### Enhanced phosphorylation of HRS upon AnnexinA1 depletion suggests that Hrs is dephosphorylated at membrane contact sites

We and others have shown that inhibition/depletion of PTP1B enhances HRS (and STAM) phosphorylation^27,28^. Since phosphorylated HRS is partly cytosolic, interaction with PTP1B would not necessarily depend on ER:endosome membrane contact. However, the enhanced HRS phosphorylation on AnnexinA1 depletion strongly suggests that HRS is dephosphorylated at membrane contacts. Notably, HRS phosphorylation was eventually lost even in the absence of Annexin A1, as previously shown in cells depleted of PTP1B^27,28^. This indicates either the presence of alternative phosphatases or that phosphorylated HRS is degraded. Consistent with the latter possibility, in cells overexpressing c-Cbl, both ubiquitylation and phosphorylation of HRS are enhanced leading to phosphorylation-dependent HRS degradation^25^. The phosphorylation state of HRS following EGF stimulation depends on the balance between phosphorylation and dephosphorylation. Although EGF stimulated HRS phosphorylation is mediated by non-receptor kinases downstream of the EGFR^20^, it should be noted that AnnexinA1 depletion and PTP1B depletion both increase the amount of EGFR on the MVE limiting membrane, which may then directly or indirectly enhance HRS phosphorylation. Enhanced HRS phosphorylation on AnnexinA1 depletion may therefore be due to a combination of increased EGF-stimulated phosphorylation and reduced PTP1B-mediated dephosphorylation.

### Effects of AnnexinA1 depletion indicate that HRS dephosphorylation is required for ILV formation

We previously showed that enhanced HRS phosphorylation on PTP1B depletion was accompanied by reduced ILV formation. Comparing the effects of AnnexinA1 depletion on ILV formation in cells expressing wild-type versus phospho-deficient HRS provided a way to distinguish between the role of phosphorylation (only possible in wild-type cells) and dephosphorylation (unnecessary in phospho-deficient cells). AnnexinA1 depletion reduced ILV formation in cells expressing wild-type HRS but ILV formation was unaffected in cells expressing HRS-Y329/334F. This strongly suggests that it is dephosphorylation of HRS that is necessary for efficient ILV formation. If HRS phosphorylation inhibits ILV formation, cells expressing phosphorylation-deficient HRS might be expected to have more ILVs than wild-type cells. However, HRS phosphorylation is rapidly lost in wild-type cells suggesting that this inhibition may be transient. Even a transient delay in ILV formation caused by sustained HRS-phosphorylation could result in prolonged EGFR engagement with downstream signalling cascades.

How might HRS phosphorylation inhibit ILV formation albeit transiently? HRS phosphorylation has been reported to promote its dissociation into the cytosol^4,21^. This would leave less HRS on the limiting membrane of the MVE to recruit other components of the sorting machinery. The increase in mean size of the coat containing phosphorylation-deficient HRS is consistent with this possibility, although the effect was modest. HRS phosphorylation could also affect its ability to bind other sorting components, including STAM, clathrin and Tsg101. Hrs binding to clathrin *in vitro* does not depend on EGF-stimulated phosphorylation^8^ but whether HRS binding to clathrin is modulated by dephosphorylation is unknown. Future studies will aim to determine the effects of Hrs phosphorylation/dephosphorylation on its interactome.

### ESCRT0 has a role in MVE:ER membrane contact site formation

Given that HRS appears to be dephosphorylated at MCSs, it is intriguing that it also has a role in MCS formation or maintenance. Our finding that STAM depletion also reduces MCS formation confirms that this is an ESCRT0-dependent role of Hrs. On the other hand, we previously found abundant MVE:ER contacts in cells depleted of the ESCRT1 component, Tsg101^31,39^, suggesting the role of ESCRT0 is distinct from the ESCRT machinery as a whole. The ER-associated protein, ORP5, has been identified in the interactome of the UIM domain of HRS^42^ raising the possibility of a direct tethering role of Hrs at MCSs. Alternatively, EGF-stimulated tyrosine phosphorylation of AnnexinA1 is required for tethering EGFR-containing MVEs to the ER^30^ and so HRS-mediated concentration of EGFR in clathrin-coated domains on the limiting membrane of MVEs could be required for AnnexinA1 phosphorylation. Arguing against this possibility is our previous demonstration that cells expressing a mutant EGFR that is negligibly ubiquitinated and cannot engage HRS, forms extensive contacts between the ER and EGFR-containing MVEs^39^. Here we show that HRS phosphorylation is not required for MVE:ER contact site formation but cells expressing phosphorylation deficient HRS showed a tendency towards more extensive contacts, suggesting a possible role of HRS phosphorylation in membrane contact maintenance/dissociation. APEX2-tagged HRS provided an opportunity to visualise the HRS-containing coat and its relationship with contact sites by conventional TEM. We have previously proposed that the HRS-containing clathrin coat would preclude the formation of membrane contacts with the ER^43^ and the distribution of the APEX2-positive coats supports this. Frequent profiles in thin section TEM and tomography show examples where the HRS-containing coat is immediately adjacent to contacts with the ER. Given the dependence of Hrs phosphorylation on PI3P-dependent association with endosomes, and the fact that clathrin recruitment is an early event in ESCRT-mediated sorting, it is likely that Hrs is phosphorylated within the clathrin coat prior to dephosphorylation at contacts formed at the edge of the coat (Figure 8D). Although we assume that both coat and contact are disassembled before ILV formation, the occasional strongly APEX2-positive ILVs indicate that HRS does not always have to dissociate from the limiting membrane of the MVE before ILV scission.

Our data re-enforce the central role of membrane contact sites in regulating membrane trafficking, in this case through promoting local dephosphorylation of a component of the sorting machinery. This opens the door for new research investigating how the contacts are regulated by this sorting machinery to co-ordinate contact disassembly with ILV formation.

## Supporting information

Video 1 for tomogram in Figure 5

Video 2 for model in Figure 5

## Acknowledgements

This work was funded by MRC (P010091/1) and Wellcome Trust (212216/Z/18/Z) grants awarded to CF and MC is recipient of a Royal Society industry fellowship, INF\R2\212031, DG was funded by a Wellcome Trust PhD studentship 105353/Z/14/Z and TB was funded by MRC career development award (MR/X020827/1). The authors gratefully acknowledge the Institute of Ophthalmology Imaging Unit for their help and support in this work.

