## Supplementary Figures for "HRS dephosphorylation at membrane contact sites promotes sorting within multivesicular endosomes"

### Supplementary Figure 1

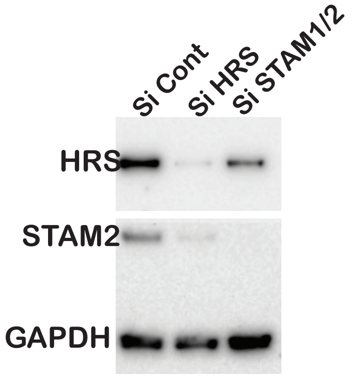


Western blot of HeLa cells treated with control siRNA or siRNAs targeting HRS or both STAMs 1 and 2 blotted for HRS, STAM2 and GAPDH as a loading control.

#### Supplementary Figure 2

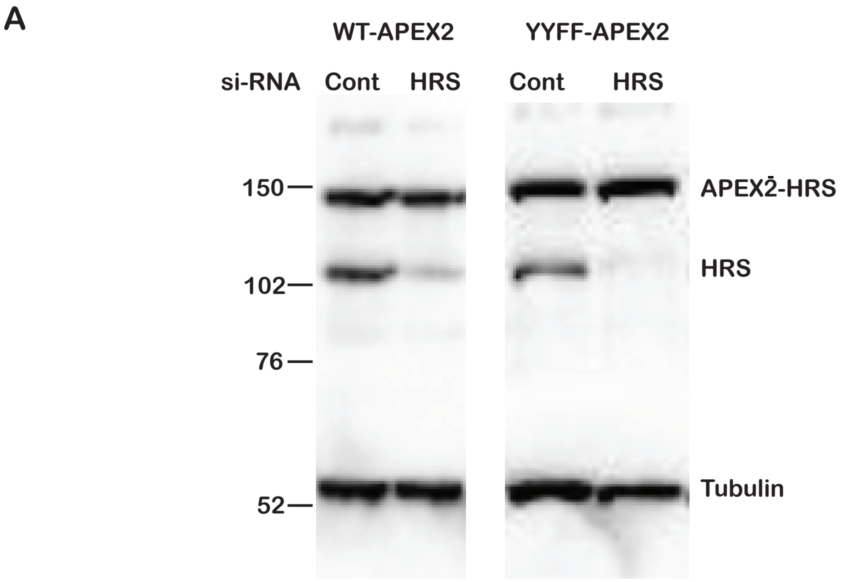


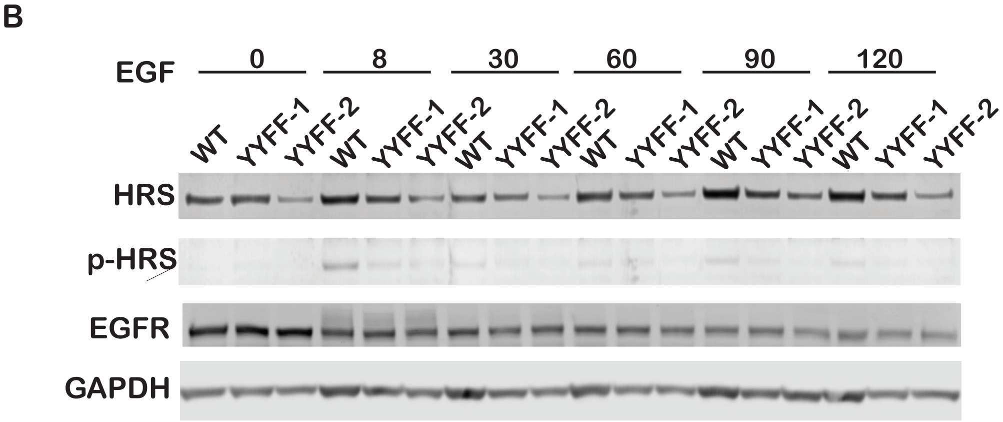


A.Western blot of Flp-In HeLa cells expressing wild-type or Y329/334F HRS were treated with control si-RNA or si-RNA targeting HRS and Western blotted with anti-total HRS and anti-tubulin antibodies. Position of molecular weight markers is indicated.

B. Western blot of hTERT-RPE cells expressing wild-type or Y329/334F HRS stimulated with EGF for the indicated times and blotted with anti-total HRS, anti-phospho Y334-HRS, anti-EGFR and anti-GAPDH antibodies

### Supplementary Figure 3

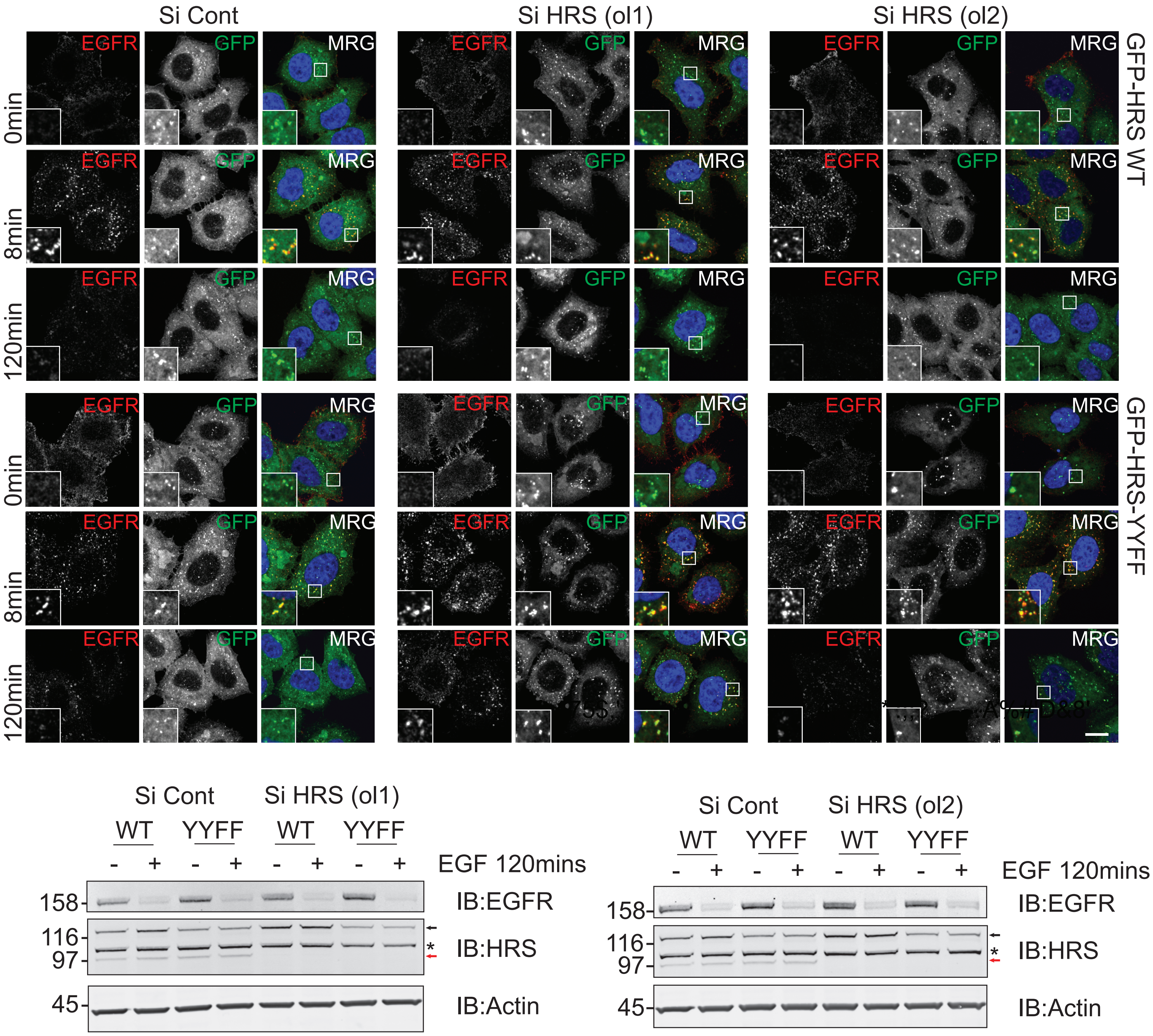


HeLaS3 stably expressing GFP-mHRS or GFP-mHRS-YYFF were depleted for endogenous HRS using two different siRNAs (ol1 and ol2) before stimulation with 20 ng/ml EGF for the indicated time. Endogenous EGFR levels were assessed by a) Immunofluorescence and b) Western blot analysis. In the HRS blot, GFP-mHRS and endogenous HRS are indicated by black and red arrows respectively and a non-specific band is indicated by *. Scale bar = 10 μm.

#### Supplementary Figure 4

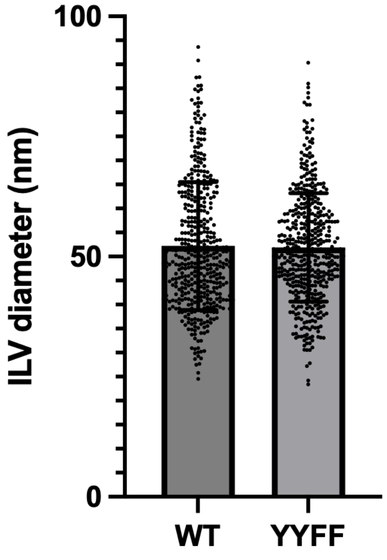


ILV diameters of flp-in HeLa cells expressing wildtype or Y329/334F HRS and stimulated with EGF and anti-EGFR-gold for 30mins. Results are means+/-SD of >100 ILVs..
